# Small proline-rich proteins (SPRRs) are epidermally produced antimicrobial proteins that defend the cutaneous barrier by direct bacterial membrane disruption

**DOI:** 10.1101/2021.09.01.458578

**Authors:** Chenlu Zhang, Zehan Hu, Abdul G. Lone, Methinee Artami, Marshall Edwards, Christos C. Zouboulis, Maggie Stein, Tamia A. Harris-Tryon

## Abstract

Human skin functions as a physical barrier, preventing the entry of foreign pathogens while also accommodating a myriad of commensal microorganisms. A key contributor to the skin landscape is the sebaceous gland. Mice devoid of sebocytes are prone to skin infection, yet our understanding of how sebocytes function in host defense is incomplete. Here we show that the small proline-rich proteins, SPRR1 and SPRR2 are bactericidal in skin. SPRR1B and SPPR2A were induced in human sebocytes by exposure to the bacterial cell wall component lipopolysaccharide (LPS). Further, LPS injected into mouse skin triggered the expression of the mouse SPRR orthologous genes, *Sprr1a* and *Sprr2a*, through stimulation of MYD88. Both mouse and human SPRR proteins displayed potent bactericidal activity against MRSA (methicillin-resistant *Staphylococcus aureus*), *Pseudomonas aeruginosa* and skin commensals. Thus, *Sprr1a^-/-^;Sprr2a^-/-^* mice are more susceptible to MRSA and *Pseudomonas aeruginosa* skin infection. Lastly, mechanistic studies demonstrate that SPRR proteins exert their bactericidal activity through binding and disruption of the bacterial membrane. Taken together, these findings provide insight into the regulation and antimicrobial function of SPRR proteins in skin and how the skin defends the host against systemic infection.

## Introduction

The skin is the human body’s largest organ, with direct contact with the external environment ^1^. As a result, the skin surface continuously encounters a diverse microbial community including bacteria, fungi, viruses and parasites ^2–4^. When host defense is impaired, skin infection results. Thus, skin and soft tissue infections pose a considerable public health threat ^5^. The majority of infections of the skin are caused by *Staphylococcus aureus* (*S. aureus*) ^6^. Additionally, *Pseudomonas aeruginosa* (*P. aeruginosa*) infections in burn patients are one of the most common causes of mortality ^7^. Adding to the challenges of treating infections posed by these pathogens has been the development of antibiotic resistant strains of bacteria such as methicillin resistant *Staphylococcus aureus* (MRSA) ^8^.

Skin antimicrobial proteins (AMPs) play an essential role in defending the host from the invasion of pathogens ^9,10^. Mammalian antimicrobial proteins are evolutionarily ancient immune effectors that rapidly kill bacteria by targeting bacterial cell wall or cell membrane structures ^11,12^. Several distinct antimicrobial protein families, such as β-defensins, cathelicidins, resistin and S100 proteins, have been identified and characterized in skin ^13,14^. However, we still have a limited understanding of the arsenal of antimicrobial proteins expressed by the skin, the regulatory networks that control the expression of AMPs, and how AMPs function to protect mammalian skin surfaces. Even less is known about the contribution that skin appendages make to host defense.

Sebaceous glands (SGs) are specialized epithelial cells that cover the entire skin surface except the palms and soles. SGs excrete a lipid-rich and waxy substance called sebum to the skin surface ^15–17^. SGs are believed to contribute to the antimicrobial functions of the skin ^13^, yet few current studies have examined the role of SGs in skin host defense. Here we show the impact of the bacterial cell wall component lipopolysaccharide (LPS) on human sebocytes and demonstrate that bacterial lipoproteins stimulate the expression of members of the small proline-rich protein (SPRR) family. SPRR proteins were originally identified as markers of terminal differentiation and function as substrates of transglutaminase in the crosslinked cornified envelope present at the skin surface ^18,19^. In this study, we demonstrate that SPRR proteins function as antimicrobial proteins. *Sprr1a^−/−^;Sprr2a^−/−^* mice are more susceptible to MRSA and *P. aeruginosa* skin infection, revealing that SPRR proteins protects against pathogenic bacterial infections of the skin. We also show mechanistically that the bactericidal activity of SPRR proteins is mediated through the binding and disruption of bacterial membranes. Taken together our findings support a novel antimicrobial function of SPRR proteins in skin.

## Results

### SPRR proteins are induced by LPS in human sebocytes

As a first step towards understanding the role of the sebaceous gland in skin host defense, we performed whole transcriptome RNA-sequencing to compare transcript abundances in human immortalized sebaceous gland cells (SZ95) treated with lipopolysaccharide (LPS) to untreated sebocytes (Fig 1A). LPS, a lipoprotein which coats the surface of Gram-negative bacteria, had broad impacts on gene expression in human sebocytes, including the increased expression of inflammatory cytokines, chemokines and antimicrobial proteins (Fig 1A). Notably, three members of the small proline rich family of proteins (SPRR) were markedly upregulated in sebocytes after LPS treatment (Fig 1A). To confirm that the expression of SPRR genes were induced by LPS, we used quantitative reverse transcription PCR (qRT-PCR) to analyze the change of the human *SPRR1B, SPRR2A, SPRR2D* transcript abundance in SZ95 cells. Consistent with our RNA-seq results, LPS-treated SZ95 cells displayed significant increase in the relative expression of *SPRR* family genes compared to vehicle treated cells (Fig 1B). Additionally, dose response and time course experiments revealed that 1μg LPS treatment for 16 hours is optimal for induction of *SPRR* family genes expression (fig S1). Consistent with the qRT-PCR results, SPRR proteins expression also increased after LPS treatment in SZ95 cells (Fig 1C, 1D). LPS is a pattern associated molecular pattern (PAMP) known to trigger gene transcription through the pattern recognition receptor, Toll-like receptor (TLR) 4 ^20^. We therefore tested a panel of PAMPs to see if other foreign stimuli would trigger the expression of the *SPRR* genes in human sebocytes. Interestingly, two other lipoproteins, Pam3CSK4 and FSL-1, stimulate the expression of *SPRR1B, SPRR2A*, and *SPRR2D* genes in human sebocytes (fig S2), indicating that sebocytes respond to bacterial lipoproteins by expressing SPRR1 and SPRR2 proteins.

**Fig. 1.**
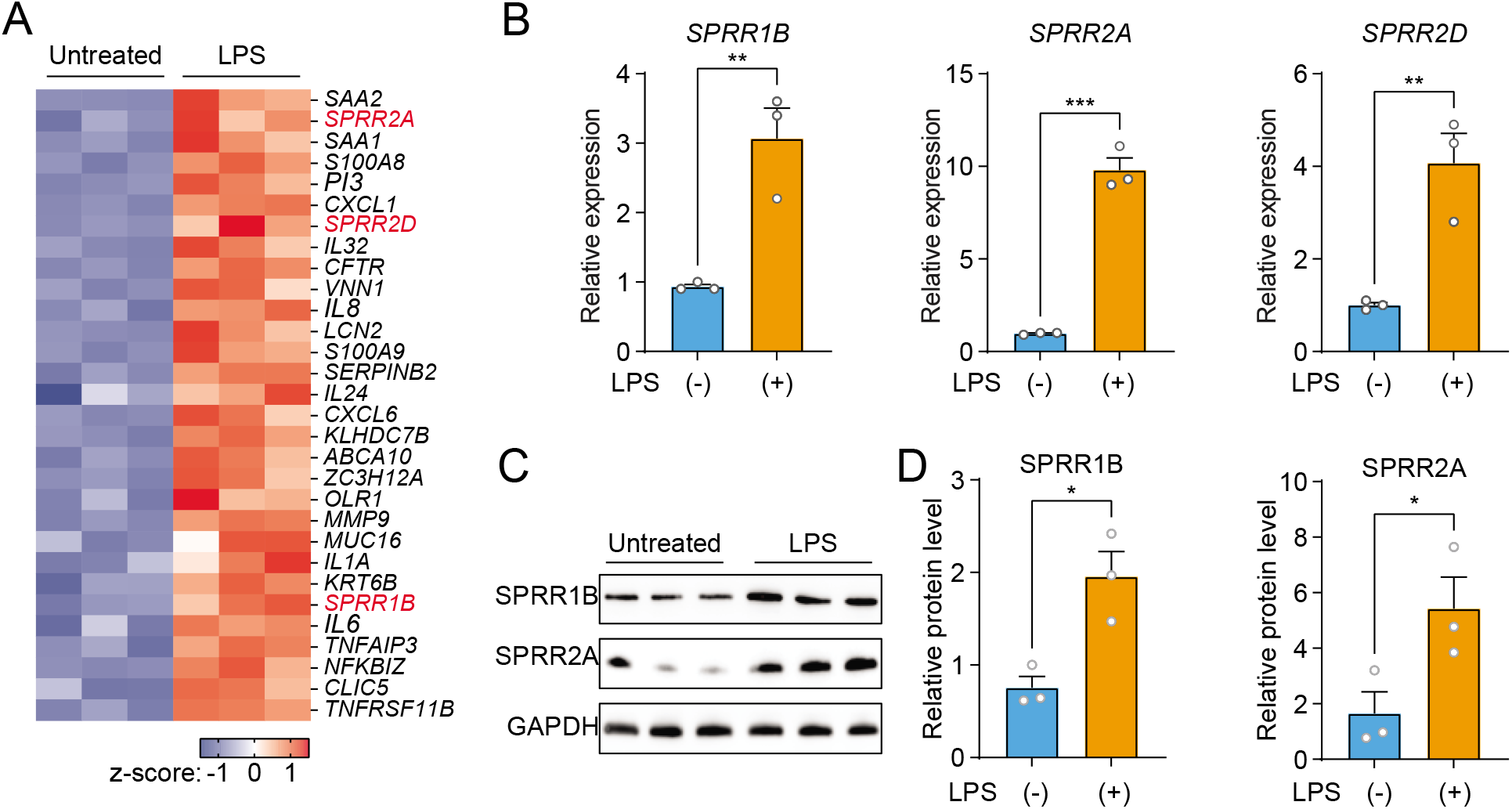
The expression of *SPRR* family genes are up-regulated by lipopolysaccharide (LPS) in human sebaceous gland cells. (A) Schematic of facial skin derived immortalized human SZ95 sebaceous gland cells treated with LPS for RNA sequencing. (B) Volcano plot with significantly up-regulated genes (fold change>1.4 and adjusted *p* value<0.01) highlighted in red (up regulated in LPS treated). (C) Heat map of significantly up-regulated genes, represented as Z-scored RPKM (reads per kilo base per million reads). *SPRR1B, SPRR2A* and *SPRR2D* are highlighted in red. (D) Reverse transcription-quantitative polymerase chain reaction (RT-qPCR) analysis of *SPRR1B, SPRR2A* and *SPRR2D* transcript in the vehicle and LPS treated human SZ95 sebocytes. (E) Western blot analysis of SPRR1B and SPRR2A was performed on vehicle or LPS treated SZ95 cells. GAPDH was used as the loading control. (F) Quantification of Western blot in (E). Means ± SEM are plotted, \**P*<0.05, \*\**P*<0.01, \*\*\**P*<0.001 was determined by unpaired t-test.

Next, we decided to further explore whether heat-inactivated bacteria could induce the expression of SPRR genes in sebocytes. We used qRT-PCR to analyze the change of the human *SPRR2A* transcript abundance in SZ95 cells after various heat-inactivated bacteria treatment. Interestingly, only the Gram-negative bacteria tested could trigger SPRR2A expression (fig S3). Taken together, these data establish that bacterial lipoproteins triggers the expression of *SPRR* genes in human sebocytes.

### SPRR proteins are upregulated by the injection of LPS in mouse skin

Next, we sought to examine whether commensal skin microbiota colonization could induce the expression of *Sprr* family genes *in vivo. Sprr1b* is not expressed in mouse skin, so we tested the ability of the microbiota to stimulate the expression of the mouse orthologs, *Sprr1a* and *Sprr2*a. In contrast to what we observed in human sebocytes in culture, *Sprr* gene expression was similar between germ free and conventional mice (Fig 2A), indicating that bacterial colonization alone is insufficient to trigger increased expression of *Sprr* genes in mouse skin. As sebaceous glands cells are located deeper in the dermis, we next chose to test whether intradermal injection of LPS – mimicking wounding and infection conditions — could stimulate *Sprr1a* and *Sprr2a* expression in mouse skin. Indeed, qRT-PCR analysis revealed that intradermal injection of LPS stimulates the expression of *Sprr1a* and *Sprr2a* in mouse skin (Fig 2B). Canonically, LPS stimulates gene expression through activation of TLR4 and the TLR signaling adaptor MYD88. Thus, LPS injected into the skin of *Myd88^-/-^* mice did not trigger increased expression of the *SPRR* genes (Fig 2B), indicating that LPS induces SPRR protein expression through the TLR-MYD88 signaling pathway. Consistent with these data, both live and heat-inactivated *Escherichia coli* (*E. coli*) bacteria were able to induce SPRR expression, suggesting that stimulation of *Sprr1a* and *Sprr2a* in mouse skin does not depend on the presence of bacterial metabolites (Fig 2C). As has been shown by others ^21^, we also found that *Sprr1a* and *Sprr2a* transcript abundance is locally induced by crosshatch skin wounding in mice (Fig 2D).

**Fig. 2.**
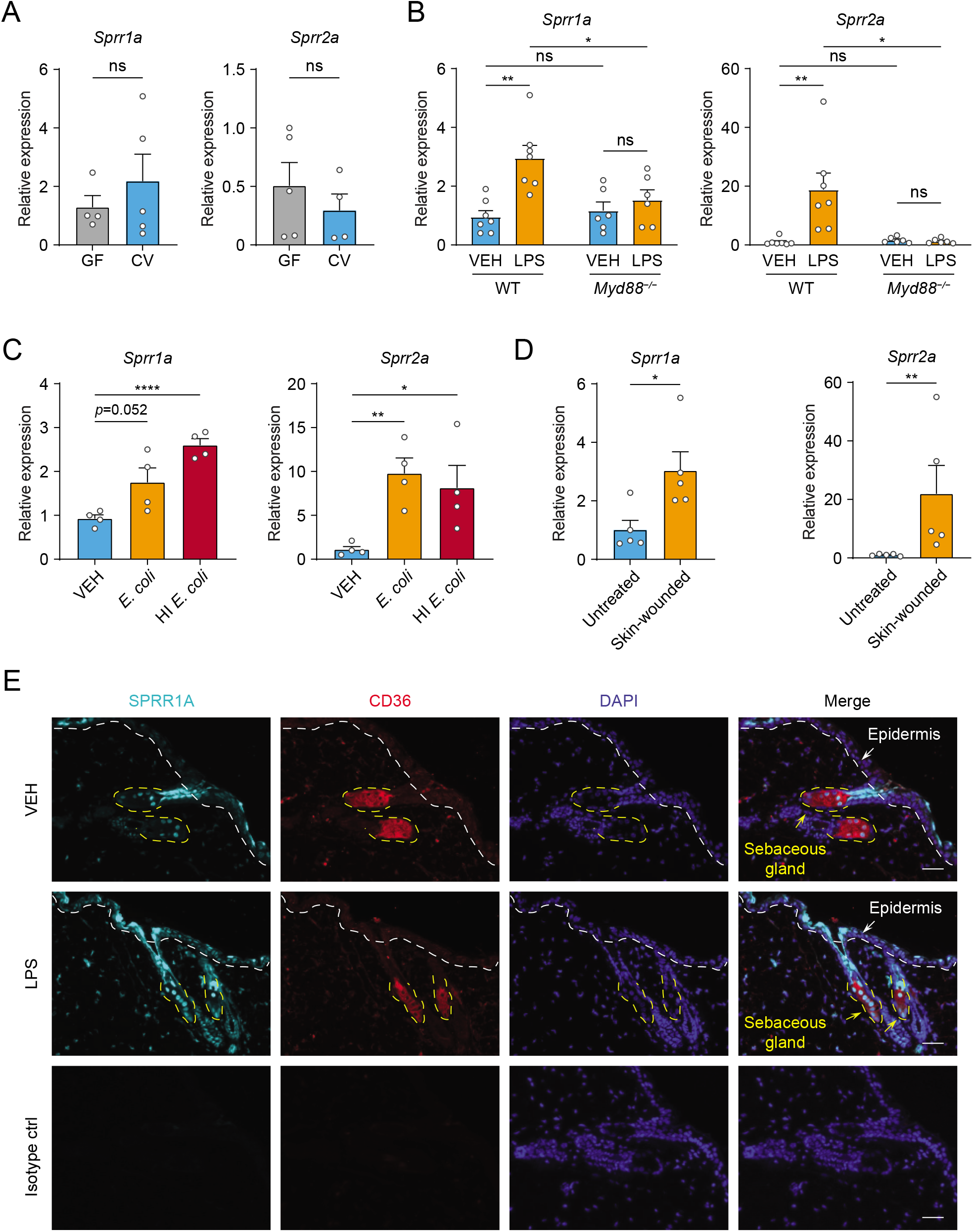
The expression of *SPRR* family genes are up regulated by LPS in mice. (A-D) qRT-PCR analysis of *Sprr1a* and *Sprr2a* gene expression in mouse dorsal skin tissue. (A) Germ-free mice (GF) compared to conventionally raised mice (CV). (B) PBS treated mouse skin compared to LPS intradermal injection mouse skin in WT or MYD88^−/−^ mouse. (C) Mouse intradermally-injected by vehicle (PBS), *E.coli* or heat-inactivated (HI) E.coli. (D) Untreated mouse skin compared to wounded skin abraded in a crosshatch pattern by a 15-blade scalpel. Means ± SEM are plotted. *P < 0.05; **P < 0.01; ****P < 0.0001ns, not significant by unpaired t test. (E) Immunofluorescence staining of SPRR1A expression in mice skin. CD36 was used as marker of sebocyte cells. Nuclei are stained with DAPI (Blue). Epidermis and sebaceous gland indicated with an arrow. White dashed line separates the epidermis and dermis. Yellow dashed line indicates the outline of SG. Scale bar, 50 μm.

Next, to investigate the expression pattern of SPRR proteins in mouse skin, we performed immunofluorescence experiment using a commercial antibody specific to SPRR1A. SPRR1A is expressed in the sebaceous gland and non-specifically at the junction of the sebaceous gland and hair follicle (Fig 2E). This suggests SPRR proteins expressed in the sebocyte may be secreted to the hair canal and are present at the skin surface. Since our immunofluorescence images and prior studies in keratinocytes indicated that SPRR1A and SPRR2A are expressed by keratinocytes ^22,23^, we sought to study whether LPS could stimulate SPRRs expression in keratinocyte cells. We first made use of hTert cells originally derived from human foreskin keratinocytes and immortalized by expression of human telomerase and mouse Cdk4 and treated the hTert cells with LPS for 24 hours. The qRT-PCR results showed that LPS cannot induce the expression of *SPRR* family genes in human keratinocytes (fig S4). To further confirm this result, we isolated primary mouse keratinocytes from neonatal mice, and cultured them under both low and high calcium condition. Under low calcium conditions the keratinocyte cell can proliferate but won’t differentiate into a stratified layer. High calcium did initiate the differentiation process and triggered increased expression of both mouse *Sprr1a* and *Sprr2a* (fig S5). However, LPS did not stimulate the expression of *Sprr* genes in mouse keratinocyte under either condition (fig S5). Taken together, these data show that SPRR proteins are unregulated by LPS in human sebocytes in culture and by LPS dermal injection in mouse skin.

### SPRR proteins are bactericidal against skin pathogens and commensals

The SPRR family proteins have highly conserved amino acid sequences with numerous cysteine and proline repeats. The antimicrobial properties of proline rich proteins have not been described in humans. However, in insects and lower vertebrates, proteins rich in cysteines and prolines have been shown to kill microbes ^24,25^. Based on these findings, we hypothesized that the SPRR proteins might function in cutaneous host defense as antimicrobial proteins. To test the antimicrobial ability of SPRR proteins, we produced recombinant human SPRR1B, SPRR2A and mouse SPRR1A protein in the baculovirus insect cell expression system and further purified to homogeneity using size exclusion chromatography (fig S6). We then tested the antimicrobial function of SPRR proteins against skin commensals and pathogens *in vitro*. When bacteria were exposed to SPRR proteins, we observed a marked dose-dependent reduction in the viability of *Staphylococcus epidermidis* (*S. epidermidis*), Methicillin Resistant *Staphylococcus aureus* (MRSA), and *P. aeruginosa* (Fig 3A). with is a more than 90% decline in bacterial viability after 2-hour exposure to 2.5 μM of SPRR proteins (Fig 3B). In contrast, SPRR proteins had no impact on the viability of the prototypical Gram-negative bacteria *Escherichia coli* (fig S7). Lastly, to visualize morphological changes of *P. aeruginosa* after exposure to SPRR proteins, we used transmission electron microscopy and observed distinct bacterial cell membrane damage and cytoplasmic leakage (Fig 3C).

**Fig. 3.**
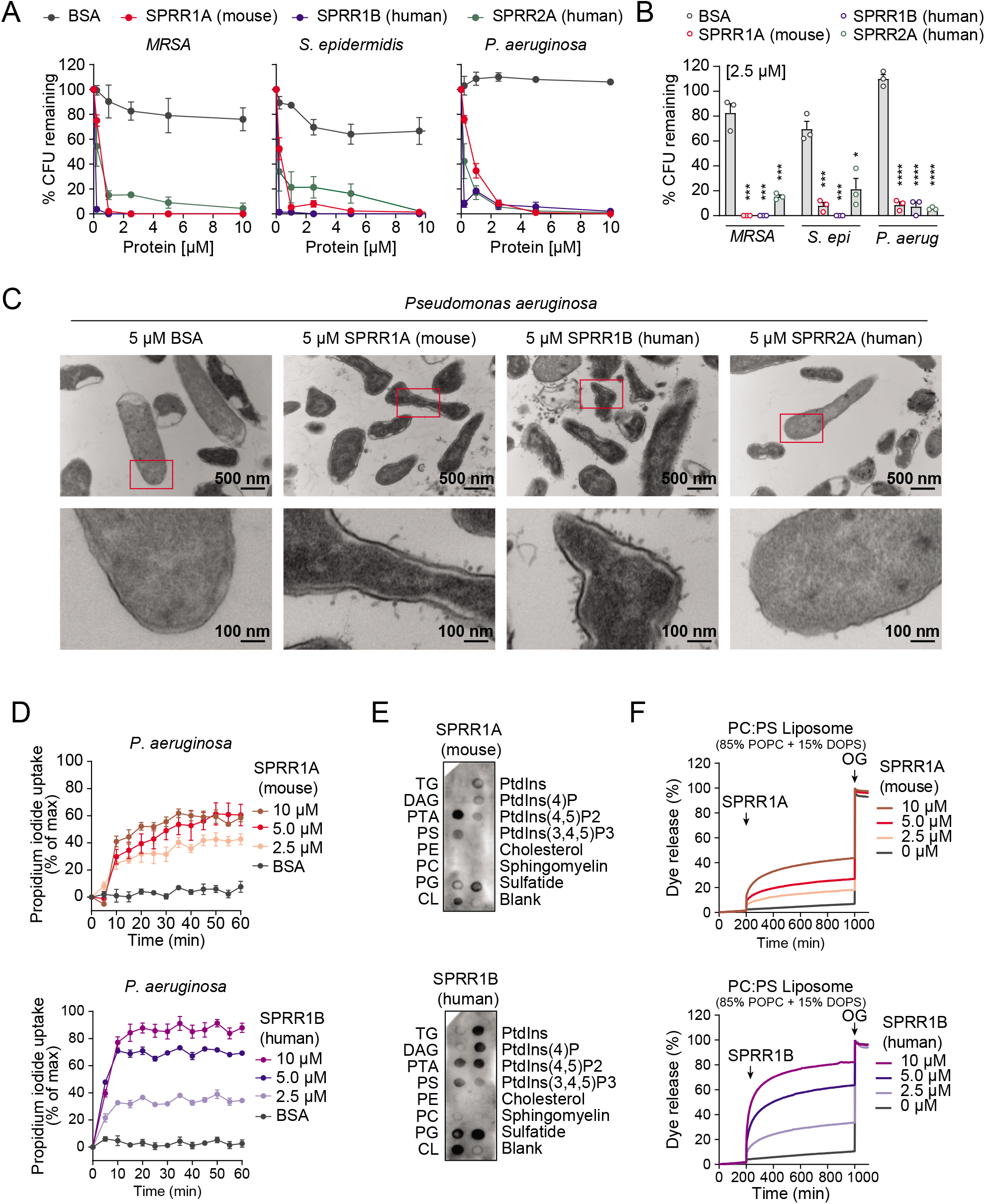
SPRR family proteins exert bactericidal activity against various skin commensal and pathogenic bacteria by membrane disruption. (A) Increasing concentrations of purified recombinant SPRR proteins were added to mid-logarithmic phase *MRSA, S. epidermidis, P.aeruginosa* for 2 hours and surviving bacteria were quantified by dilution plating. (B) 2.5 μM of SPRR proteins was added to midlogarithmic phase bacteria for 2 hours and surviving bacteria were quantified by dilution plating. Remaining Colony forming units (CFUs) are expressed as a percentage of untreated bacteria control. (C) Transmission electron microscopy of *P. aeruginosa* after incubation with 5 μM purified recombinant SPRR proteins. BSA was used as negative control. Examples of cell surface damage and cytoplasmic leakage are indicated with red rectangular box. Upper scale bars, 500 nm. Lower scale bars, 100 nm. (D) Propidium iodide (PI) uptake by *P. aeruginosa* in the presence of increasing concentrations of mouse SPRR1A and human SPRR1B protein. (E) Membranes displaying various lipids were incubated with 1μg/mL SPRR proteins, and detected with specific antibody. (F) Carboxyfluorescein (CF)-loaded liposomes were treated with increasing concentrations of mouse SPRR1A and human SPRR1B protein, and dye efflux was monitored over time and was expressed as a percentage of total efflux in the presence of the detergent octyl glucoside (OG). All assays were performed in triplicate. Means ± SEM are plotted. *P < 0.05; ***P < 0.001; ****P < 0.0001; ns, not significant by two-tailed t-test.

Since SPRR proteins exhibit similar spectrum of bactericidal activity, we chose to use human SPRR1B and mouse SPRR1A protein to further delineate the mechanism underlying its bactericidal activity. We first used the propidium iodide (PI) uptake assay to assess the capacity of SPRR proteins to permeabilize bacterial membranes. Human SPRR1B and mouse SPRR1A proteins promoted the dose-dependent uptake of the membrane-impermeant small molecule dye, PI, by *P. aeruginosa* (Fig 3D). Next, we examined whether SPRR proteins can directly bind lipid molecules by incubating proteins with lipid strips dotted with different lipids. Both human SPRR1B and mouse SPRR1A bind to lipids bearing negatively charged lipid head groups, but not to neutral lipids (Fig 3E). These data are consistent with the idea that most bacterial cell membranes are negatively charged and thus susceptible to SPRR1 binding. Lastly, to determine whether SPRR proteins disrupt bacterial membrane, we incubated human SPRR1B and mouse SPRR1A protein with liposomes loaded with the fluorescent dye carboxyfluorescein (CF, ~10 Å Stokes diameter) and quantified dye release after exposure. Phosphatidylcholine (PC)/ phosphatidylserine (PS) liposomes are composed of 85% of the neutral lipid PC and 15% of the negatively charged lipid PS —similar to that of bacterial membranes. Both human SPRR1B and mouse SPRR1A protein induced rapid dye efflux from negatively charged PC/PS liposomes in a dose-dependent manner (Fig. 3F). These findings confirm that SPRR proteins exert bactericidal activity through membrane disruption. Taken together, these results suggest that SPRR proteins are bactericidal and compromise the membrane of specific skin bacteria including the pathogen *P. aeruginosa*.

### SPRR proteins limit MRSA and *P. aeruginosa* infection

Given that SPRR1A and SPRR2A protein exhibited bactericidal activity against a panel of skin pathogens *in vitro* (Figure 3), we predicted that the removal of these proteins might limit MRSA and *P. aeruginosa* skin infection *in vivo*. To test this hypothesis, we created mice lacking SPRR1A and SPRR2A. We used a CRISPR/Cas9-mediated in vitro fertilization method to delete the entire mouse *Sprr1a* locus of the *Sprr2a^−/−^* mice ^26^ and verified that SPRR1A and SPRR2A are absent in *Sprr1a^−/−^;Sprr2a^−/−^* mice skin at both the transcript and protein level (fig S8). *Sprr1a^−/−^;Sprr2a^−/−^* mice were healthy and showed normal skin barrier with no signs of immune infiltration and no significant change in transepidermal water loss (TEWL) (fig S9).

To test the *Sprr1a^−/−^;Sprr2a^−/−^* mice for susceptibility to infection, we inoculated the skin of wild-type and *Sprr1a^−/−^;Sprr2a^−/−^* mice with 1 x 10^6^ CFU of a bioluminescent strain of MRSA by topical application. The luminescence was monitored daily with serial photography and quantified for 3 consecutive days (Fig 4A). Mice devoid of SPRR1A and SPRR2A expression had greater bacterial burdens in the first 3 days of MRSA infection (Fig 4B, 4C). Next, we aimed to assess the susceptibility of *Sprr1a^−/−^;Sprr2a^−/−^* mice to *P. aeruginosa* skin infection. Since damaged or burned skin has increased risk of *P. aeruginosa* infection ^7^, we first introduced skin wounding in mice by superficial crosshatching method, and then inoculated 1 x 10^6^ CFU of a bioluminescent strain of *P. aeruginosa* at the wounding site. Intravital imaging and quantification revealed that *Sprr1a^−/−^;Sprr2a^−/−^* mice are more susceptible to *P. aeruginosa* skin infection (Fig 4D-4F). Collectively, we confirm that SPRR proteins limit MRSA and *P. aeruginosa* infection in mice skin.

**Fig. 4.**
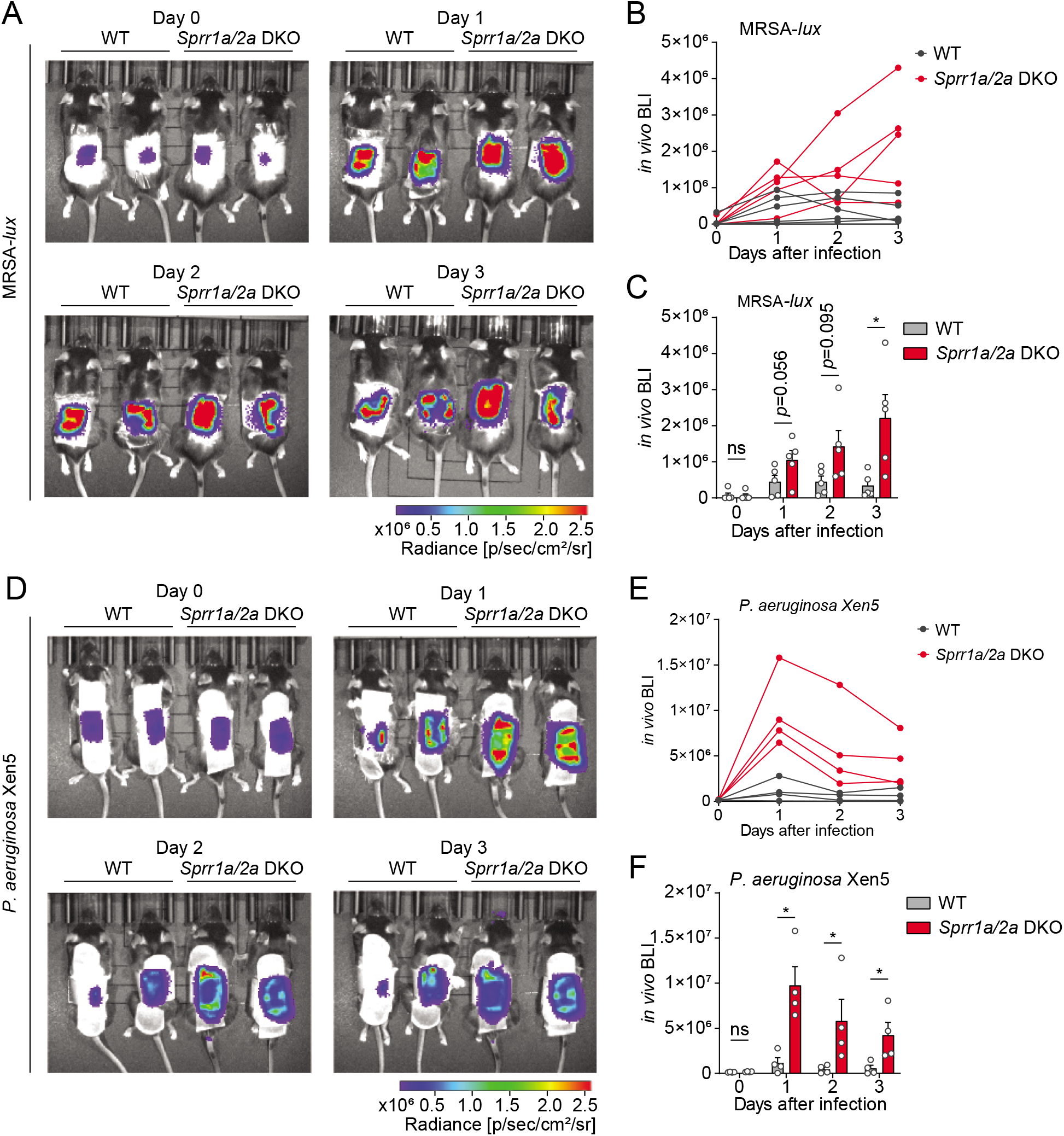
SPRR family proteins protect against skin MRSA and *P. aerugonisa* infection. (A-C) WT and *Sprr1a^−/−^;Sprr2a^−/−^* mice were epicutaneously challenged with MRSA (1 x 10^6^ CFU) on the shaved dorsal skin for 3 consecutive days. (A) Representative *in vivo* bioluminescent imaging (BLI) photographs from Day 0 to Day 3. Quantification of MRSA total flux (photons/s) for each mouse (B) or plotted as Means ± SEM (C). (D-F) WT and *Sprr1a^−/−^;Sprr2a^−/−^* mice were superficially abraded in a crosshatch pattern by a 15-blade scalpel to introduce skin wounding. One day post crosshatch wounding, WT and *Sprr1a^−/−^;Sprr2a^−/−^* mice were topically applied with 1 x 10^6^ CFU *P. aerugonisa* on the back skin for 3 days. (D) Representative in vivo bioluminescent imaging (BLI) signals from Day 0 to Day 3. Quantification of *P. aerugonisa* total flux (photons/s) for each mouse (E) or plotted as Means ± SEM (F). *p < 0.05, ns, not significant by unpaired t-test.

## Discussion

Skin is in direct contact with the microbe-filled outer world. Thus, defending the host from invasion by pathogens like *S. aureus* and *P. aeruginosa* is one of the chief functions of the skin ^6,27^. In this study we have identified SPRR1 and SPRR2 proteins as a previously uncharacterized group of antibacterial proteins expressed by keratinocytes and sebaceous gland cells, which can rapidly kill pathogens through membrane disruption and limit bacterial skin infection.

Several earlier studies have shown that SPRR proteins are upregulated in the GI tract, urinary tract and the airway after exposure to stress and other inflammatory stimuli ^28^. Further, Hu et al. recently revealed that SPRR2A has antimicrobial actions in the gastrointestinal tract during helminth infection ^26^. Our data highlight a previously unappreciated role of SPRR family proteins in skin immune defense, demonstrating that both SPRR1 and SPRR2 are bactericidal proteins induced in sebocytes by the bacterial cell wall component LPS (Fig 1). The major role of sebaceous glands in mammals is to produce sebum, a mixture of non-polar lipids and proteins required for normal skin ecology ^29^. Sebum secretion can also act as a delivery system for AMPs ^30^. Though beneficial to the host, AMPs can damage mammalian membranes, so confining expression of AMPs to the non-viable parts of skin through sebum delivery is optimal. Sebum also excretes AMPs to the surface within an acidic milieu, which is often required for AMP activity ^31^.

Interestingly, our data also show that bacterial colonization of germ-free mice did not induce expression of SPRR proteins *in vivo* (Fig 2). Further, LPS was not able to stimulate the production of SPRR proteins in mouse keratinocytes (fig S4-5). Taken together, our findings suggest that deeper penetration of bacterial stimuli may be required for MYD88 mediated SPRR protein production in skin. These finding are distinct from what has been observed in the gastrointestinal tract, where colonization of germ-free mice is sufficient to stimulate SPRR2A expression ^32^. As SPRR family proteins also function as cross-linking proteins in terminal differentiation of keratinocytes and formation of the cornified cell envelope (Candi et al., 2005), there are likely other regulatory networks that control SPRR proteins expression in skin in response to wounding and other stimuli. Additional studies with sebaceous gland-specific deletion of TLRs and MYD88 will be required to confirm that stimulation of SPRR proteins by LPS requires interaction with TLRs on sebocytes.

In this study, we also reveal that SPRR family proteins have potent bactericidal activity against *Staphylococcal* species and the gram-negative bacteria *P. aeruginosa* (Fig 3A-C), with effective inhibition at low micromolar concentrations. Further, our biochemical data show that mouse and human SPRR1 proteins interact with negatively charged lipid membranes, suggesting broad spectrum antimicrobial capability of these proteins (Fig 3D-F). Additionally, skin devoid of SPRR1 and SPRR2 is more susceptible to skin infection by MRSA and *Pseudomonas aeruginosa*, both pathogens that cause skin infections requiring hospitalization (Fig 4). Treatments for these infections are currently limited. SPRR proteins might be promising alternatives to traditional antibiotics used to cure MRSA or *P. aeruginosa* infection.

Human SPRR proteins are 6–18 kDa in size and comprise four subclasses (SPRR1, SPRR2, SPRR3, and SPRR4) with a similar structural organization. Among them, two SPRR1 and seven SPRR2 proteins are characterized with a much higher homogeneity, as they contain a similar consensus repeat sequence ^18^. Our findings that SPRR family proteins have antimicrobial function may have implications for other proline-rich proteins expressed at other mucosal sites. Altogether, these findings expand our current understanding of the molecules involved in cutaneous host defense and provide insight into how the sebaceous gland contributes to the fight against skin infection.

## Materials and Methods

### Mice

All animal protocols were approved by the Institutional Animal Care and Use Committees of the University of Texas (UT) Southwestern Medical Center. Age- and sex-matched mice 8-12 weeks old were used for all experiments. Conventionally raised C57BL/6 wild type (WT), Myd88^-/^ and *Sprr1a^−/−^;Sprr2a^−/−^* on C57BL/6 background were bred and maintained in the specific pathogen free (SPF) barrier facility at the UT Southwestern Medical Center on standard chow diet. The generation of *Sprr1a^−/−^;Sprr2a^−/−^* mice is described below. Germ-free (GF) C57BL/6 mice were bred and maintained in flexible film vinyl isolators in the gnotobiotic mouse facility at UT Southwestern with autoclaved diet (Hooper Lab Diet 6F5KAI, Lab Diet, St. Louis, MO) and autoclaved nanopore water. All mice were housed under a 12 hour light-dark cycle, and mice were randomly assigned to treatment groups.

*Sprr1a^−/−^;Sprr2a^−/−^* mice were generated using CRISPR/Cas9-mediated *in vitro* fertilization with a guide RNA targeting regions upstream and downstream of the *Sprr1a* locus on the basis of *Sprr2a^−/−^* mice (Figure S8). *Sprr2a^−/−^* mice were obtained with the permission of Dr. Lora Hooper ^26^. Guide RNAs were injected into fertilized female *Sprr2a^−/−^* mice embryos by the UT Southwestern Transgenic Core facility. The resulting litters were screened by genomic sequencing to detect the deletion of *Sprr1a* and *Sprr2a* locus. Mice harboring the deleted allele were backcrossed with WT to homozygosity.

### Bacterial strains

Bacteria were grown in species-specific growth media: *Escherichia coli* (ATCC; ATC-PTA-7555) and *Citrobacter rodentium* (ATCC51459) were grown in Luria Broth. *Pseudomonas aeruginosa* (ATCC27853), bioluminescent *Pseudomonas aeruginosa* strain Xen05 (Perkin Elmer 119228) and *Enterococcus faecalis* (ATCC29212) were grown in Brain Heart Infusion Broth (BD Biosciences). Methicillin resistant *Staphylococcus aureus* (ATCC25923), bioluminescent MRSA strain SAP430 *lux* ^33^, *Staphylococcus epidermidis* (clinical isolate), *Staphylococcus xylosus* (clinical isolate) were grown in Tryptic Soy Broth.

### Sebocyte cell culture and treatments

SZ95 cells are an immortalized human sebocyte cell line generated from the face of an 87-year-old female and transformed with Simian virus 40. These cells were previously obtained from Cristos Zouboulis ^14^. SZ95 sebocytes were maintained in Sebomed Basal Medium (Fischer Scientific NC9711618) supplemented with 10% fetal bovine serum (GeminiBio 100-106), 5 ng/ml human epidermal growth factor (ThermoFisher PHG0313) and 1% Antibiotic-Antimycotic (Gibco 15240062). Cells were cultured in 5% CO_2_ incubator at 37°C. Cells were stimulated with 1μg/ml LPS (Sigma L4524). 16 hours post-stimulation cells were harvested and analyzed as described below. For TLR agonists treatment, reagents in human TLR1-9 Agonist kit (InvivoGen, tlrl-kit1hw) were diluted to working concentration in PBS. 100ng/ml Pam3CSK4, 1 x 10^8^ cells/ml HKLM, 500ng/ml Poly (I:C), 500ng/ml Flagellin, 100ng/ml FSL-1 and 1μg/ml Imiquimod were used for stimulation of SZ95 cells. For heat-inactivated bacteria treatment, bacteria were grown to mid-logarithmic phase, spun down, washed and resuspended in PBS. Bacteria were heat-inactivated by incubation at 95 °C for 20 minutes. Then 1 x10^8^ cells/ml were used for each treatment.

### Immunofluorescence microscopy

Mouse skin samples were fixed in formalin and embedded in paraffin by the UT Southwestern histology core. Samples were deparaffined with xylene followed by rehydration with decreasing concentrations of ethanol. Boiling in 10mM sodium citrate buffer with 0.2% tween for 15 minutes for antigen retrieval. Slides were blocked with 10% FBS, 1% BSA and 1% Triton X-100 in PBS, and then incubated with primary antibodies against mouse SPRR1A (Thermo Fisher PA5-26062), mouse CD36 (R&D AF2519), rabbit IgG isotype control (Thermo Fisher 02-6102) or goat IgG isotype control (Thermo Fisher 02-6202) using 1:100 dilutions at 4°C overnight. Secondary antibodies Alex Fluor 594 or Alexa Fluor 647 (Thermo Fisher) were diluted 1:350 in blocking buffer and applied to slides for 1 hour at room temperature in the dark. Slides were then washed with PBST (PBS with 0.2% tween) and mounted with DAPI Fluoromount-G (Southern Biotechnology 0100-20). Images were captured using an Echo Revolve 4 microscope.

### Quantitative real-time PCR

RNA was extracted from cells or mouse skin using the RNAeasy Plus universal kit (Qiagen 73404). RNA was quantified by absorbance at 260 nm, and its purity was evaluated by the ratios of absorbance at 260/280 nm. 2μg of RNA was used for cDNA synthesis (Thermofischer 4368814, High Capacity cDNA reverse transcription kit). Quantitative real-time PCR was performed using PowerUp SYBR Green Gene Expression Assays (Thermofischer A25741) and a QuantStudio 7 Flex Real-Time PCR System (Applied Biosystems). Relative expression values were calculated using the comparative Ct (ΔΔCt) method, and transcript abundances were normalized to GAPDH transcript abundance. The primer sequences are shown in table S1.

### Whole transcriptome sequencing and data analysis

RNA was extracted from SZ95 sebocyte cells using the RNAeasy Plus universal kit (Qiagen 73404). RNA quality was assessed using Agilent 2100 Bioanalyzer. Truseq RNA sample preparation kit v2 (Illumina) was used for the preparation of sequencing libraries. Sequencing was performed on an Illumina HiSeq 2500 for signal end 50 bp length reads. Sequence reads were mapped against the hg19 genome using TopHat. For each gene, read counts were computed using HTSeq-count and analyzed for differential expression using DESeq2.

### Western blot

SZ95 cells were harvested by applying 200 μL of diluted 1x sample buffer (Thermo Fisher 39000) directly to a 6-well plate, scraping down the cell sample to disrupt the membranes, then boiling for 15 minutes before loading. Equal amounts of protein were loaded onto a 4-20% gradient SDS-PAGE and transferred to a PVDF membrane. After blocking with 5% milk in TBST, the membranes were incubated with anti-SPRR1B antibody (Thermo Fisher PA5-26062), anti-SPRR2A (Abcam ab125385) or anti-GAPDH (Abcam ab181602) at 4°C overnight. Membranes were then incubated with anti-rabbit secondary antibodies conjugated with HRP (Abcam). Membranes were visualized using a Bio-Rad ChemiDoc Touch system and bands were quantified by Image Lab software.

### Keratinocyte cell line culture and treatments

hTERT (ATCC CRL-4048) and primary mouse keratinocyte cells were cultured in KSFM medium (Invitrogen, 37010022) supplemented with 0.05 mM CaCl_2_ (Sigma, C7902), 0.05 μg/mL hydrocortisone (Sigma, H0888), 5 ng/mL EGF (Invitrogen, 10450-013), 7.5 μg/mL bovine pituitary extract (Invitrogen, 13028-014), 0.5 μg/mL insulin (Sigma, I9278), 100 U/mL penicillin, 100 μg/mL streptomycin, and 25 μg/mL of amphotericin B (Invitrogen, 10450-013). Mouse primary keratinocytes were isolated through dispase digestion. Before digestion, the subcutaneous fat was first removed from the mouse skin. Skin tissue from 3-5 days neonatal mice was floated on 1 U/ml dispase (Corning, 354235) in HBSS (Gibco, 14170) for 16 h at 4°C with the dermis side down. The next day, the skin was placed in a new dish with the epidermis side down, and the epidermis was peeled and placed into a new dish with HBSS. After being washed, the cells were collected into a new 15 mL tube. The epidermis was cut into small pieces, resuspended in HBSS, gently pipetted up and down several times, and then combined with the cells in the 15 mL tube. The cell solution was filtered with a 70 μm cell strainer, centrifuged at 1,000 rpm for 4 minutes, and resuspended in complete KSFM medium. The cells were gently washed once and seeded in culture dishes with complete KSFM medium. The culture dishes were precoated with collagen (Advanced Biomatrix, 5005-B). Keratinocytes from each mouse were seeded in one 10 cm dish, and fresh complete KSFM medium was supplied after 24 hours. After 3-4 days, the primary keratinocytes reached approximately 80% confluence under normal culture conditions in a 5% CO_2_, 37°C incubator. For keratinocyte differentiation, cells were treated with 1 mM CaCl_2_ for 2 days. For treatment of keratinocyte cells, 1 μg/mL LPS (Sigma L4524) were supplemented in the KSFM complete medium. 16 hours post-stimulation cells were harvested for qRT-PCR or Western blot analysis.

### Intradermal injection of mice

The mouse dorsal hair was removed by shaving (Andis ProClip), followed by depilatory cream (Nair™) one day before injection. LPS (Sigma L4524) was dissolved in PBS and further diluted to a concentration of 1 mg/mL. For intradermal injection of LPS, each mouse was injected with 50 μL of LPS solution to the back skin. For intradermal injection of bacteria, various bacteria were grown in species-specific media to mid-logarithmic phase, spun down, washed and resuspended in PBS to a concentration of 1 x10^8^ cells/mL. Bacteria were heat-inactivated by incubation at 95 °C for 20 minutes. 100 μL of live or heat-inactivated bacteria was injected to the mouse back skin. The site of injection was circled with permanent marker. After 8 hours, mice were sacrificed and the injection site skin was analyzed.

### Protein expression and purification

*SPRR* genes containing a C-terminal 6xHis tag were cloned into a pFastBac1 vector and heterologously expressed in Sf9 cells (Thermo Fisher). One liter of cells (2.5×10^6^ cells/mL) was infected with 10 mL baculovirus at 28°C. Cells were cultured in suspension and harvested 48 hours after infection. Harvested cells were resuspended in the buffer containing 25 mM Tris-HCl pH 8.0, 150 mM NaCl and 1 mM Phenylmethylsulfonyl fluoride (PMSF), and lysed by sonication. The mixture was pelleted by centrifugation at 10,000*g* for 30 minute and the supernatant was loaded onto a Ni^2+^ metal affinity column (Qiagen). The column was washed with buffer containing 30 mM imidazole and the protein was eluted in buffer containing 300 mM imidazole. The eluate was concentrated in a 3K cutoff Amicon Ultra centrifugal device (Millipore) and further purified by size exclusion chromatography on a Superdex 75™ 10/300 GL column (GE Healthcare Life Sciences) in standard assay buffer (10 mM MES, pH 6.0 and 25 mM NaCl).

### Bacterial killing assays

Bacteria were grown in species-specific growth media as described above. 10 mL bacterial cultures were grown to mid-logarithmic phase and then pelleted and washed twice in assay buffer (10 mM MES, pH 6.0, 25 mM NaCl). Approximately 5 x10^6^ cells/mL bacteria were then incubated at 37°C for 2 hours in assay buffer with varying concentrations of recombinant SPRR protein or BSA (Gemini 700-106P). Surviving colony-forming units (CFU) were quantified by dilution plating onto agar plates and calculated as a percentage of the remaining colonies in the assay buffer only control sample.

### Lipid strip assay

Membrane lipid strips (Echelon, P-6002) were used following the manufacturer’s protocol. Briefly, the lipid strips were blocked with blocking buffer (10 mM MES, pH 6.0, 25 mM NaCl, 2% BSA, and 0.05% Tween-20) for 1 hour at room temperature. Purified recombinant SPRR proteins were diluted to 1 μg/mL in blocking buffer and incubated with the lipid strip overnight at 4°C. After three washes with washing buffer (10 mM MES, pH 6.0, 25 mM NaCl and 0.05% Tween-20), the lipid strip was sequentially incubated with anti-SPRR1B antibody and HRP-conjugated secondary antibody. Dots were detected with ECL reagent (Bio-Rad, 1705060) using a Bio-Rad ChemiDoc system.

### Electron microscopy

For electron microscopy of bacteria, 10 mL *P. aeruginosa* cultures were grown to mid-logarithmic phase and then pelleted and washed in 10 mL standard assay buffer (10 mM MES pH 6.0 and 25 mM NaCl). Bacteria were then resuspended in 1 mL standard assay buffer. Purified recombinant SPRR proteins was added at a final concentration of 5 μM to 300 μL of resuspended bacteria and incubated at 37°C for 2 hours. Bacteria were centrifuged for 10 minutes at 16,000 g, resuspended in cross-linking reagent (4% paraformaldehyde and 4% glutaraldehyde in 0.1 M sodium phosphate buffer, pH 7.4), and incubated overnight at 4°C. After three washes, bacterial pellets were embedded in 3% agarose, sliced into small blocks (1 mm^3^), and fixed with 1% osmium tetroxide and 0.8% potassium ferricyanide in 0.1 M sodium phosphate buffer for 1.5 hours at room temperature. Cells were rinsed with water then stained with 1% uranyl acetate in water for 1 hour. Cells were dehydrated with increasing concentrations of ethanol, transitioned into propylene oxide, infiltrated with Embed-812 resin, and polymerized overnight in a 60°C oven. Blocks were sectioned with a diamond knife (Diatome) on Leica Ultracut 7 ultramicrotome (Leica Microsystems), collected onto copper grids, and post-stained with 2% aqueous uranyl acetate and lead citrate. Images were captured on a Tecnai G2 spirit transmission electron microscope (Thermo Fisher) equipped with a LaB6 source using a voltage of 120 kV.

### Liposome disruption assay

Unilamellar liposomes were prepared using lipids from Avanti. 85% 1-palmitoyl-2-oleoyl-*sn*-glycero-3-phosphocholine or POPC (AVANTI PC #850457C) and 15% 1,2-dioleoyl-*sn*-glycero-3-phospho-L-serine or DOPS (AVANTI PS #840035C) dissolved in chloroform were mixed in specific molar ratios in glass tubes and then dried under a stream of N_2_, followed by drying under vacuum overnight to ensure complete removal of organic solvents. Dried lipids were then combined with 5(6)-carboxyfluorescein (CF) (Sigma #21877) and vortexed for 5 minutes. Lipids were transferred to 2 mL cryotubes and subjected to five freeze thaw cycles in liquid N_2_ and stored at −80 °C freezer. Lipids were then thawed and passed through a 100 nm pore membrane using a mini-extruder kit (Avanti Polar Lipids #610000) and purified on a PD-10 column to remove excess dye. Prepared liposomes were diluted in standard assay buffer (10 mM MES, pH 6.0, 25 mM NaCl) to a working concentration of 100 μM. QuantaMaster 300 fluorometer (Photon Technology International) was used to monitor fluorescence. SPRR protein was added to the system at increasing concentrations. At the end time point, 1% v/v n-octylglucoside detergent (OG, Anatrace #O311) was added to completely disrupt the liposomes. Fluorescence was measured over time in seconds and as a percentage of total CF dye released by the detergent OG.

### Dye uptake assay

Bacterial cultures were grown to mid-logarithmic phase and then pelleted and washed in standard assay buffer (10 mM MES, pH 6.0 and 25 mM NaCl). Bacteria were then diluted to approximately 5 x 10^8^ cells/mL in standard assay buffer containing 5.5 μg/mL Propidium iodide (PI) (Thermo Fisher, P3566). Then bacterial samples (90 μL each well) were added to black 96-well Costar plates (Fisher, 07-200-567) and placed into a Spectramax plate reader (Molecular Devices) that was pre-equilibrated to 37°C for 10 minutes. After an initial reading, 10 μL of recombinant purified SPRR proteins at varying concentrations or BSA were added and fluorescence outputs (excitation, 535 nm; emission, 617 nm) were measured every 5 minutes for 1 hour. PI uptake activity was measured against the maximum fluorescence output from the positive control added with 0.05% SDS.

### Skin infections

The dorsal hair was removed from C57BL6 *Sprr1a^−/−^;Sprr2a^−/−^* or WT male mice by shaving (Andis ProClip), followed by depilatory cream (Nair). For MRSA infection, one day after shaving, bioluminescent MRSA strain SAP430-*lux* were grown to mid-logarithmic phase in Tryptic Soy Broth and then pelleted and washed twice in PBS. Approximately 1 x 10^6^ CFU bacteria in 100 μL PBS were placed on rectangular gauze. For *P. aeruginosa-* infection, after 24 hours, the dorsal skin was superficially abraded in a crosshatch pattern by a 15-blade scalpel (Fine Scientific Tools 10115-10) to introduce skin wounding. One day post-wounding, the bioluminescent *P. aeruginosa* strain Xen05 (Perkin Elmer 119228) were grown in Brain Heart Infusion Broth (BD Biosciences) and around 1 x 10^6^ CFU bacteria in 100 μL PBS were placed on a gauze rectangle. The gauze was applied to the dorsal skin of *Sprr1a^−/−^;Sprr2a^−/−^* or WT mice and held in place with 2 tegaderm (3M 9505W) and a Band-Aid Sheer Strips (BAND-AID) for 3 days. The luminescent was monitored daily with serial photography and quantification using a IVIS Lumina3 imager machine.

### TEWL measurement

Transepidermal water loss (TEWL) of mice dorsal skin was measured using Vapometer (Delfin Technologies) according to manufacturer instructions.

### Data availability

RNA-seq data (Figures 1B, 1C) has been submitted to the Gene Expression Omnibus with an accession number: GSE182756

### Statistical analysis

Statistical details of experiments can be found in the figure legends, including how significance was defined and the statistical methods used. Data represent mean ± standard error of the mean. Formal randomization techniques were not used; however, mice were allocated to experiments randomly and samples were processed in an arbitrary order. Mouse skin samples that were determined to be in the anagen hair cycle were excluded. All statistical analyses were performed with GraphPad Prism software. To assess the statistical significance of the difference between two treatments, we used two-tailed Student’s t-tests. To assess the statistical significance of differences between more than two treatments, we used one-way ANOVAs.

## Acknowledgements

We would like to thank Dr. Lora Hooper for helpful discussion and sharing *Sprr2a^−/−^* mice strain. We thank Dr. Richard Wang for sharing the hTert immortalized human keratinocyte cell line. We thank Dr. Zheng Kuang for his help with analysis of RNA-sequencing data. This work was supported by a Harold Amos Award through the Robert Wood Johnson Foundation, National Institutes of Health K08 award, a UT Southwestern Disease Oriented Clinical Scholars Program award, and a Burroughs Wellcome Fund Career Award for Medical Scientists.

## Author Contributions

C.Z. designed and performed most experiments. Z.H. performed protein expression and purification and liposome assays in fig S6 and Fig 5. A.L. performed skin infection experiments in Fig 4. M.A. and M.E. provided technical assistance with the mouse experiments and assisted with bacterial killing assays in Fig 3A, 3B. M.S. completed the dose response and time course experiments in fig S1. C.C.Z revised the manuscript and is the original source of the SZ95 cells.

C.Z. and T.A.H. designed experiments, interpreted results, and wrote the manuscript.

---

**fig. S1.**
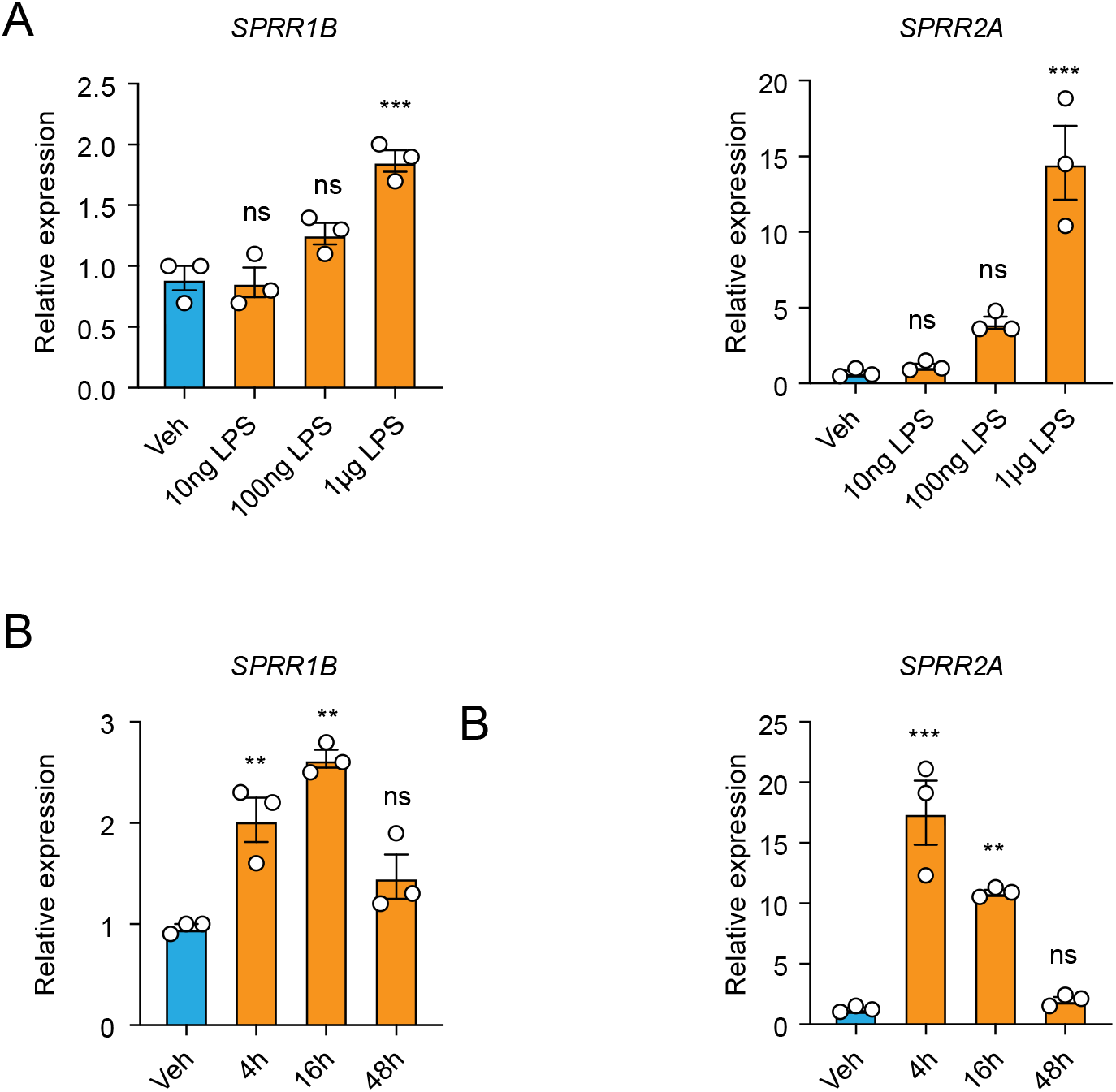
Dose response and time course analysis of LPS treatment on human sebocyte cells. qRT-PCR analysis of *SPRR1B and SPRR2A* transcript in the vehicle and various LPS treated human SZ95 sebocytes. (A) SZ95 sebocyte cells were treated with different concentration of LPS for 16 hours. (B) SZ95 cell were treated with 1 μg/mL LPS for different length of time. Means ± SEM are plotted. **P < 0.01; ***P < 0.001;ns, not significant by unpaired t-test.

**fig. S2.**
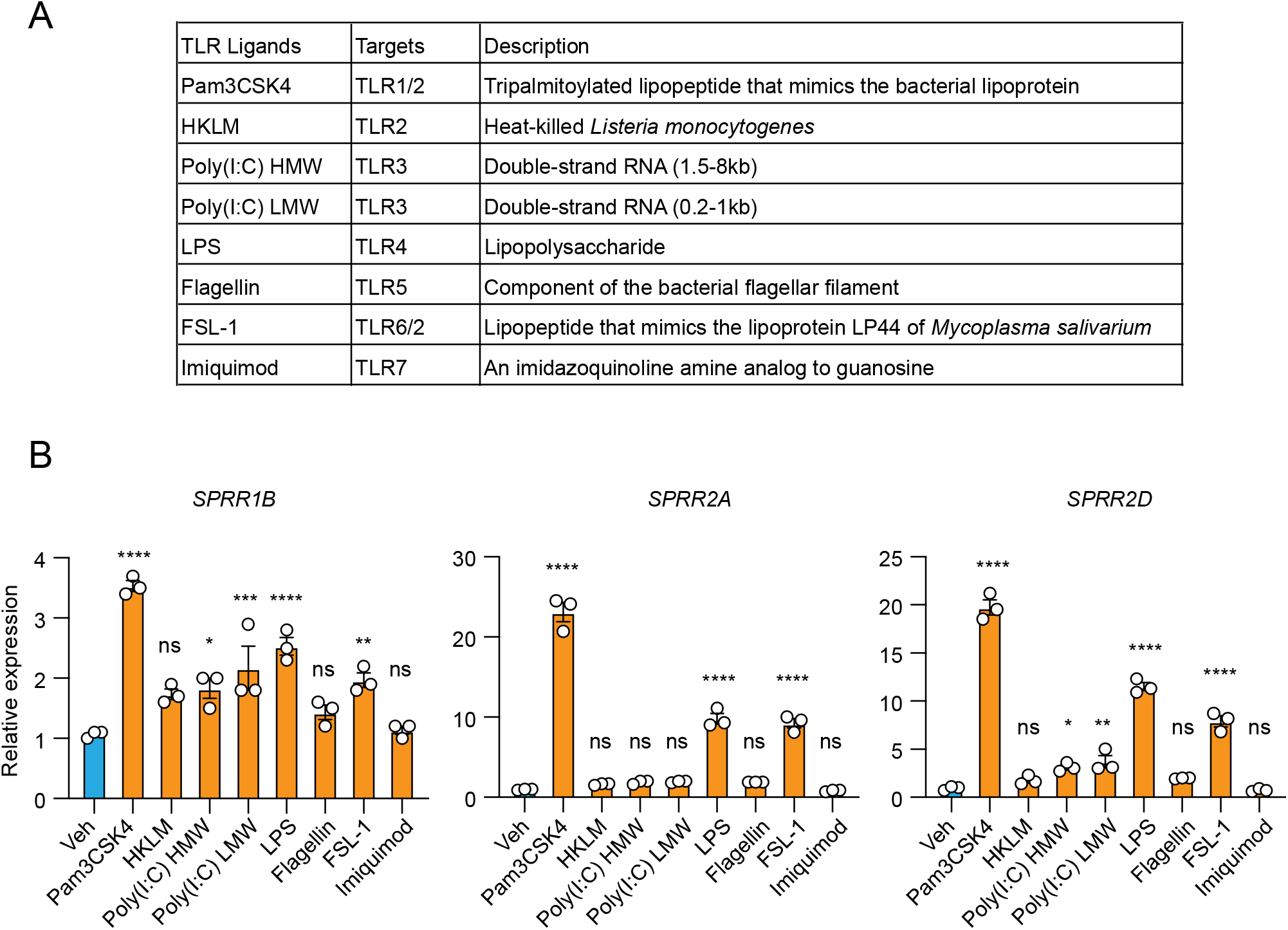
The expression of SPRR family genes are up regulated by TLR2 and TLR4 agonists in human sebaceous gland cells. (A) A list of different TLR agonists. (B) qRT-PCR analysis of *SPRR* family genes expression in the vehicle and various TLR agonists treated SZ95 cells. Means ± SEM are plotted. *P < 0.05; **P < 0.01; ***P < 0.001; ****P < 0.0001; ns, not significant by one-way ANOVAs.

**fig. S3.**
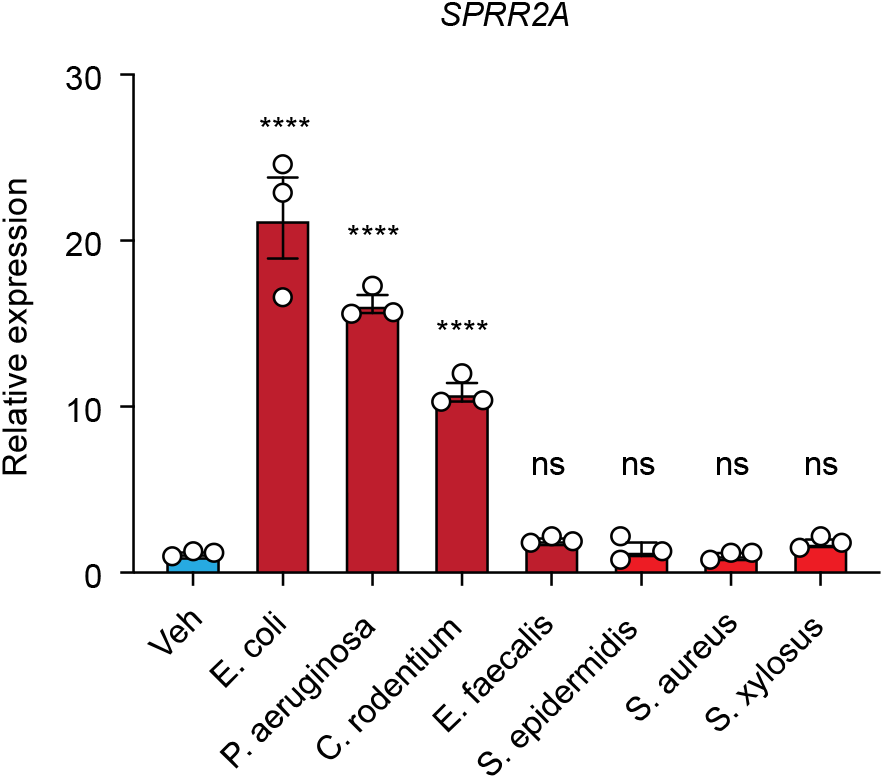
Gram-negative bacteria can trigger *Sprr2a* gene expression in human sebaceous gland cells. qRT-PCR analysis of *Sprr2a* transcript in the vehicle and various heat inactivated bacteria treated human SZ95 sebocyte cells. Means ± SEM are plotted. ****P < 0.0001; ns, not significant by one-way ANOVAs.

**fig. S4.**
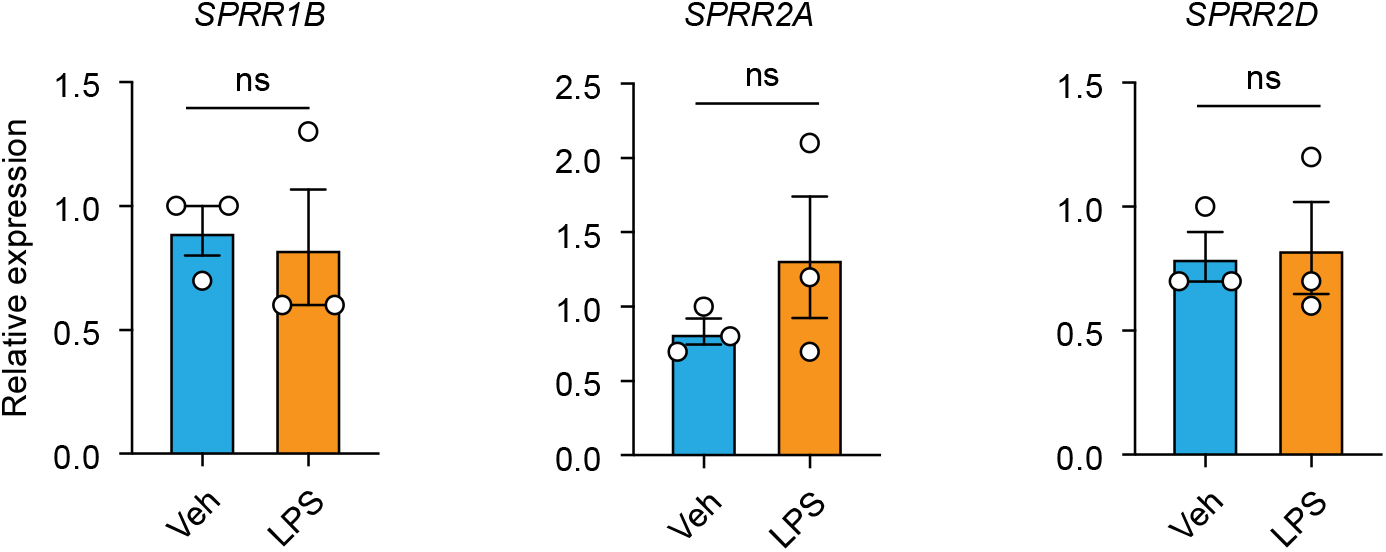
LPS cannot induce the expression of *SPRR* family genes in human keratinocyte cells. hTERT immortalized human keratinocyte cells were treated with LPS for 16 hours. qRT-PCR analysis of *SPRR1B, SPRR2A* and *SPRR2D* transcript in the vehicle and LPS treated human hTERT keratinocytes. Means ± SEM are plotted. ns, not significant by unpaired t-test.

**fig. S5.**
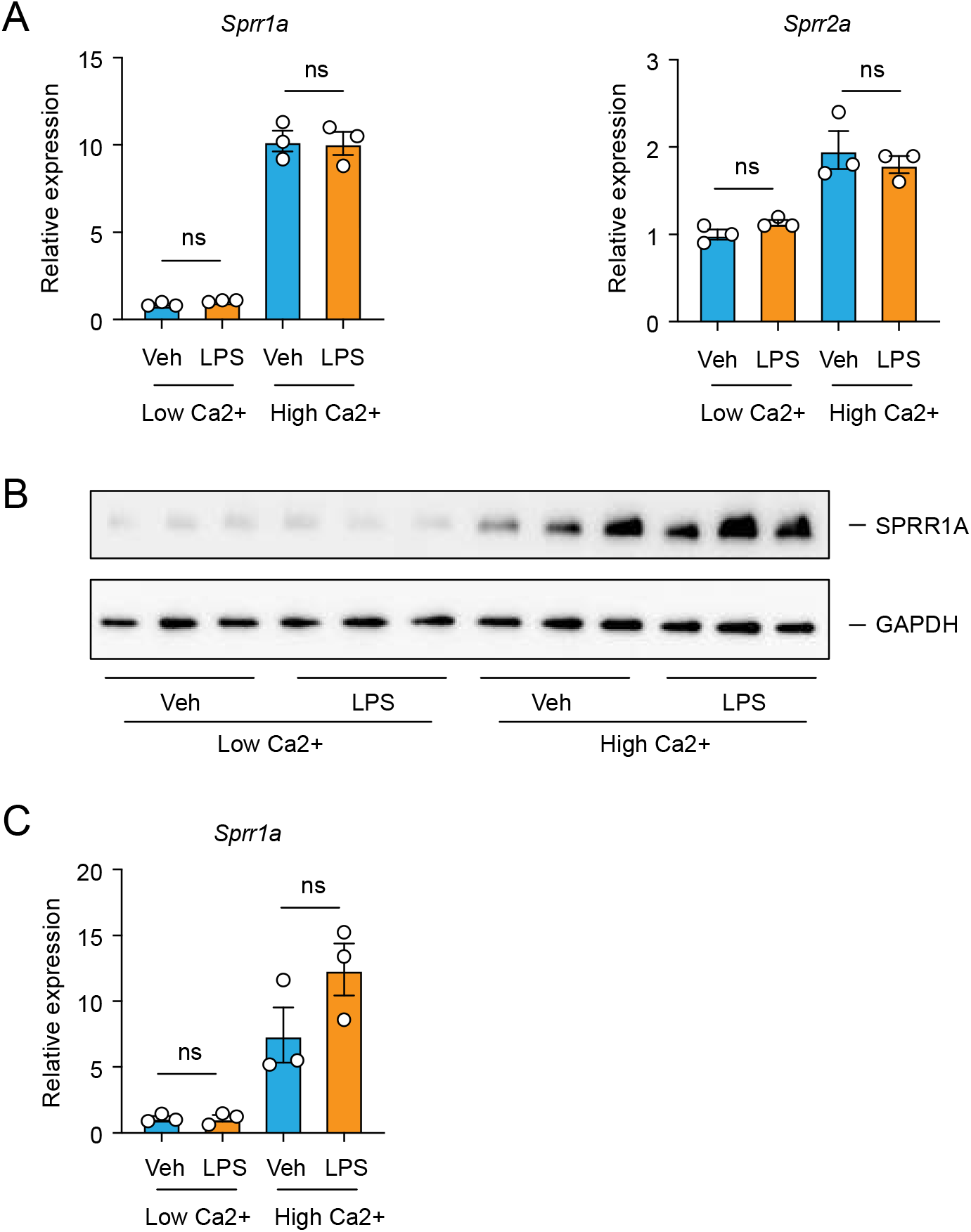
LPS cannot trigger the expression of *Sprr* family genes in primary mouse keratinocytes. Primary mouse keratinocytes were isolated from 3-5 days old neonatal mice through dispase digestion. With 0.05 mM CaCl_2_ (Low Ca^2+^), keratinocytes can proliferate but won’t differentiate into a stratified layer. Under 1 mM CaCl_2_ (High Ca^2+^) condition, keratinocytes will initiate differentiation process. (A) qRT-PCR analysis of *Sprr1a* and *Sprr2a* transcript in the vehicle and LPS treated primary mouse keratinocyte cells cultured under Low Ca^2+^ or High Ca^2+^ condition. (B) Western blot analysis of SPRR1A protein was performed on vehicle or LPS treated primary mouse keratinocytes under both Low Ca^2+^ and High Ca^2+^ condition. (C) Quantification of the Western blot in (B). Means ± SEM are plotted. *P < 0.05; ***P < 0.001 by unpaired t-test.

**fig. S6.**
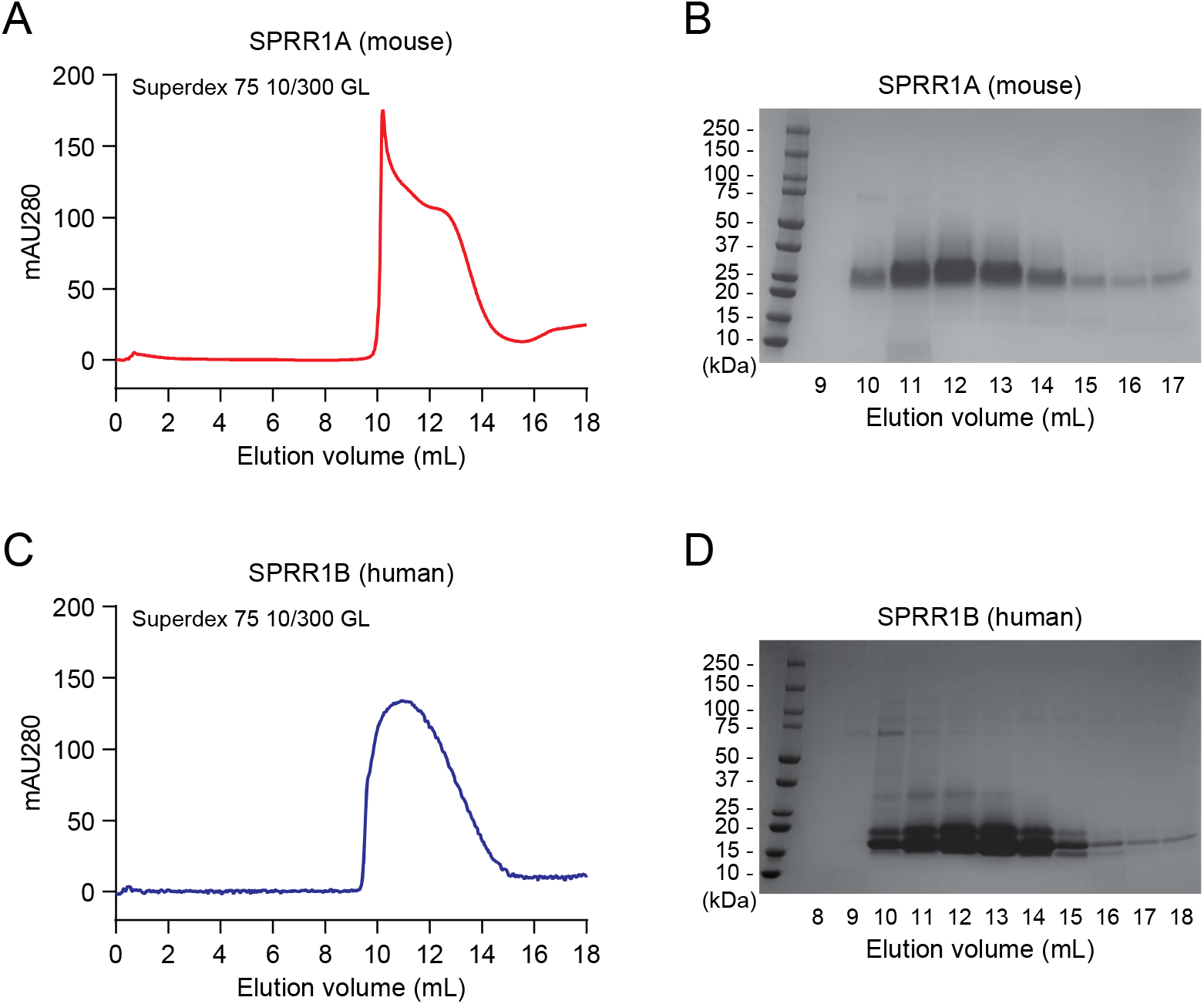
Recombinant expression and purification of SPRR proteins. (A and C) Recombinant mouse SPRR1A protein (A) and human SPRR1B protein (C) was expressed using a baculovirus expression system and purified by size-exclusion chromatography (Superdex 75 10/300 GL) ^26^. (B and D) Fractions from (A) and (C) were visualized by Coomassie blue staining following SDS-polyacrylamide gel electrophoresis (SDS-PAGE).

**fig. S7.**
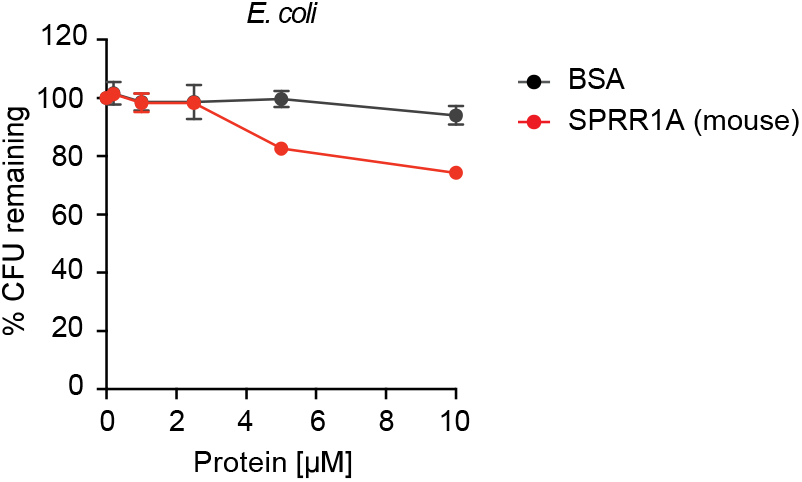
SPRR1A protein was resistant to Gram-negative bacteria *Escherichia coli*. Increasing concentrations of purified recombinant mouse SPRR1A protein was added to mid-logarithmic phase *Escherichia coli* for 2 hours and surviving bacteria were quantified by dilution plating. Remaining Colony forming units (CFUs) are expressed as a percentage of untreated bacteria control. BSA was used as negative control. Means ± SEM are plotted.

**fig. S8.**
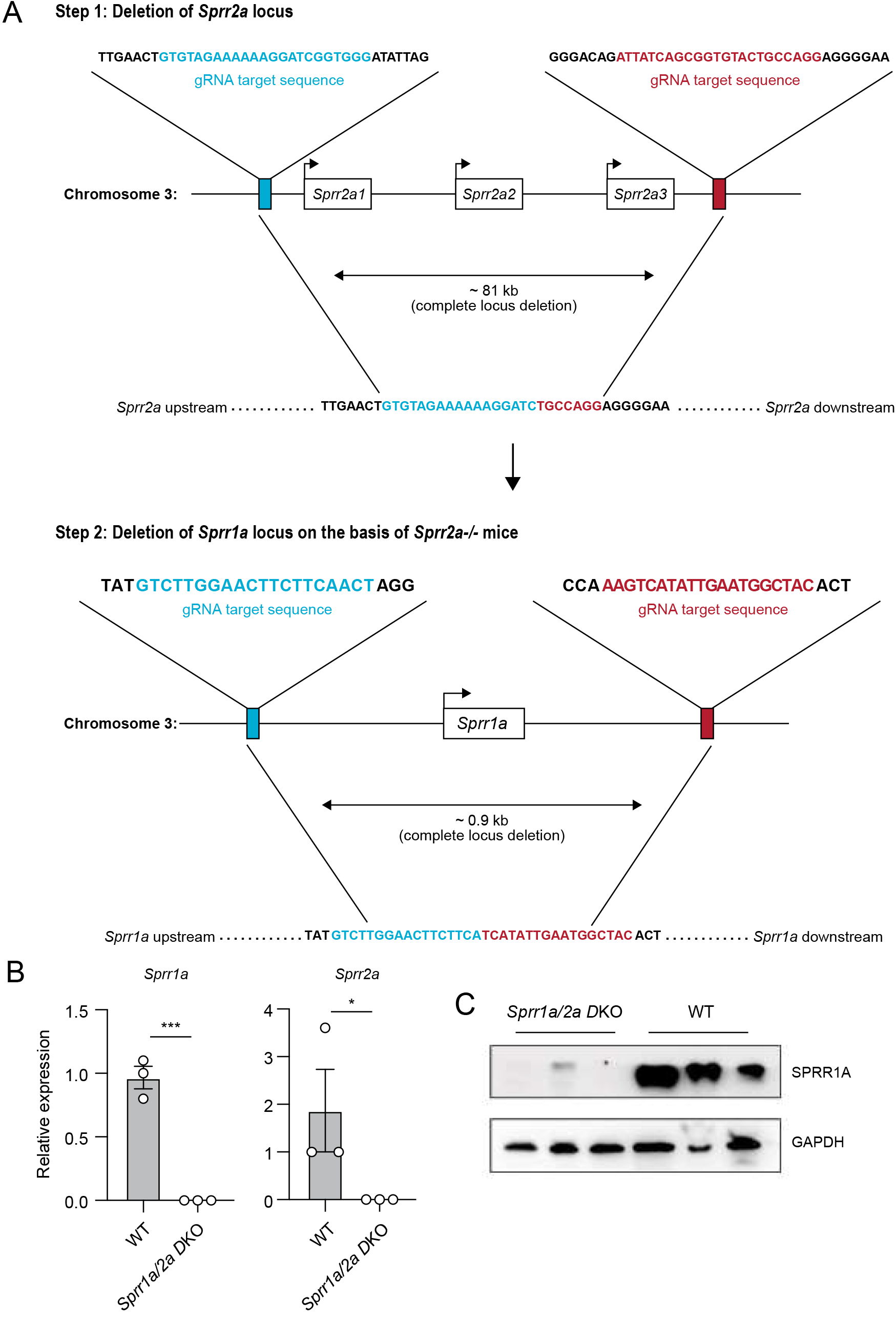
Generation and validation of *Sprr1a^−/−^;Sprr2a^−/−^* mice by CRISPR/Cas9 genomic targeting. (A) Schematic diagram of two-step strategy using CRISPR/Cas9-mediated gene targeting to delete the entire *Sprr2a* and *Sprr1a* locus. (B) qRT-PCR analysis of *Sprr1a* and *Sprr2a* expression in the skin of WT and *Sprr1a^−/−^;Sprr2a^−/−^* mice. Values were normalized to *Gapdh* expression. Means ± SEM are plotted. *P < 0.05; ***P < 0.001 by unpaired t-test. (C) Western blot analysis of SPRR1A protein in the skin of WT and *Sprr1a^−/−^;Sprr2a^−/−^* mice. GAPDH was used as the loading control.

**fig. S9.**
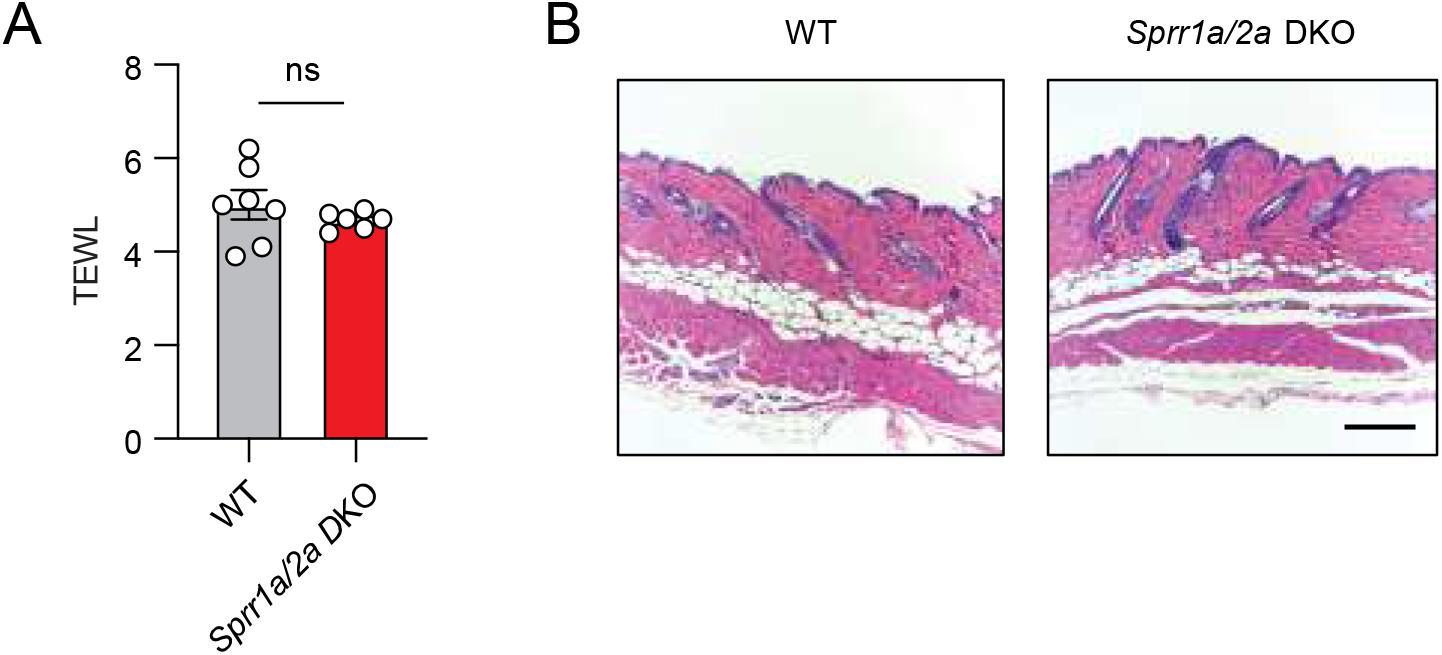
*Sprr1a^−/−^;Sprr2a^−/−^* mice does not show signs of inflammation or impaired skin barrier. (A) Transepidermal water loss (TEWL) of WT and *Sprr1a^−/−^;Sprr2a^−/−^* mice skin was analyzed. Means ± SEM is plotted, ns, not significant by unpaired t test. (B) Representative H&E staining of mouse skin from WT and *Sprr1a^−/−^;Sprr2a^−/−^*. Scale bar, 200 μm.

